# Alternative Assembly of Qβ Virus Like Particles

**DOI:** 10.1101/2022.01.23.477406

**Authors:** Vincent Shaw, Suttipun Sungsuwan, Hunter McFallBoegeman, Xuefei Huang, Xiangshu Jin

## Abstract

Qβ virus like particles (VLPs) are versatile platforms for grafting functional groups for vaccine development. The structure of Qβ VLPs at atomic detail are critical for design of more effective vaccines. While the structures of native Qβ VLPs have been determined previously, the structure of VLPs assembled from a recombinantly expressed Qβ coat protein, which are extensively used as platforms have not been studied. We sought to determine the crystal structures of VLPs assembled from recombinantly expressed Qβ coat protein of wild type and two mutants: A38K and A38K/A40C/D102C. The structures of Qβ VLPs assembled from recombinantly expressed Qβ coat proteins showed that VLPs can be assembled both in T=1 and T=3 symmetry.

## 1. Introduction

Bacteriophage Qβ is one of many small RNA viruses infecting *Escherichia coli*. Qβ Virus like particles (VLPs) are a new class of immunogenic carriers for development of vaccines (Chuang et al., 2020; Bachman et al., 2010), detailed structural knowledge of the carrier VLPs is critical for effective vaccine design. The crystal structure and cryo EM structure of the native Qβ particle were reported previously (Golmonhammadi et al., 1995; Cui et al., 2017). However, the structure of Qβ VLP assembled from recombinantly produced coat protein has not been investigated. Here we report the crystal structures of Qβ VLPs assembled from recombinantly expressed coat protein in wild-type, a mutant harboring a single mutation, A38K, and a mutant harboring triple mutation, A38K/A40C/D102C. Remarkably, these new structures show that recombinantly produced Qβ coat protein can assemble VLPs both in T=1 and T=3 symmetry.

## 2. Materials and Methods

### 2.1. Overexpression and purification of Qβ proteins

The detailed procedure for overexpression and purification of the proteins used in crystallization is essentially the same as that reported in (Sungsuwan, S. et al., 2021).

### 2.2. Crystallization and Data Collection

The crystals of wild-type Qβ VLPs, both in T=1 and T=3 symmetries, were grown by the hanging drop vapor diffusion method using 5mg/ml protein mixed with and equilibrated against a mother liquor containing 10% (wt/vol) PEG3350, 5mM cobalt chloride, 0.1M Tris, pH 8.5. Crystals were cryo-protected using the mother liquor supplemented with 25% (vol/vol) ethylene glycol. The crystals of mutant Qβ VLPs, both A38K and A38K/A40C/D102C, were grown by the hanging drop vapor diffusion method using 5mg/ml protein mixed with and equilibrated against a mother liquor containing 10% (wt/vol) PEG3350, 0.2M L-proline, 0.1M HEPES, pH 7.5. Crystals were cryo-protected using the mother liquor supplemented with 25% (vol/vol) ethylene glycol. X-ray diffraction data were collected at 100K on the LS-CAT and GM/CA-CAT beamlines at the Advanced Photon Source, Argonne National Laboratory. The diffraction data were processed using HKL2000 (Otwinowski and Minor, 1997) and scaled/merged with SCALEPACK (Otwinowski and Minor, 1997). The structure was solved by molecular replacement with PHASER (McCoy, 2007) using the coordinates of Chain A in 1QBE as the search model. Iterative model fitting and refinement were carried out using COOT (Emsley et al., 2010) and PHENIX (Adams et al., 2010), to yield the final refined structures whose statistics are detailed in Table 1. The coordinates for the four structures have been deposited to PDB with the accession codes: 7TJD (wild type T=1), 7TJE (wild type T=3), 7TJG (A38K T=1), and 7TJM (A38K/A40C/D102C T=1).

**Table 1.** Data Collection and Refinement Statistics.

| **Data collection** |  |  |  |  |
| --- | --- | --- | --- | --- |
| Qβ protein | Wild-type, T=1 | Wild type, T=3 | A38K, T=1 | A38K/A40C/D102C, T=1 |
| Beamline | APS 21-ID-D | APS 21-ID-F | APS 21-ID-F | APS 23-ID-B |
| Wavelength (Å) | 0.976 | 0.97872 | 0.97872 | 1.033167 |
| Space group | *R3* | *R3* | *I23* | *I23* |
| *Cell dimensions* |  |  |  |  |
| a, b, c (Å) | 271.9, 271.9, 166.56 | 286.6, 286.6, 662.7 | 194.1, 194.1, 194.1 | 191.7, 191.7, 191.7 |
| $\alpha$, $\beta$, $\gamma$ (°) | 90, 90, 120 | 90, 90, 120 | 90, 90, 90 | 90, 90, 90 |
| Resolution (Å) | 45.3 - 3.5 (3.62 - 3.50) | 49.5-3.6 (3.71-3.60) | 45.8 - 2.8 (2.9 - 2.8) | 30.3 - 3.9 (4.04 - 3.9) |
| Unique reflections | 57859 (5795) | 245320 (17430) | 29445 (3012) | 10727 (1074) |
| R<sub>merge</sub> | 0.101 (0.546) | 0.14 (0.58) | 0.12 (0.54) | 0.13 (0.60) |
| I/σI | 10.1 (2.5) | 9.0 (2.7) | 9.0 (2.7) | 9.0 (2.7) |
| Completeness (%) | 99.7 (99.0) | 99.9 (99.9) | 97.9 (100.0) | 98.8 (100.0) |
| Redundancy | 8.6 (3.6) | 8.3 (6.5) | 17.2 (14.8) | 14.3 (10.9) |
| Wilson B | 84.0 | 100.7 | 58.5 | 128.0 |
| **Refinement** |  |  |  |  |
| Reflections used in refinement | 57812 | 234901 | 29435 | 10703 |
| Resolution (Å) | 39.5–3.5 | 49.6-3.6 | 45.8-2.8 | 30.3-3.9 |
| Reflections used in refinement | 57812 | 234901 | 29435 | 10703 |
| Reflections used for R-free | 2903 | 12266 | 1500 | 523 |
| R<sub>work</sub> / R<sub>free</sub> | 0.195 /0.244 | 0.278/0.328 | 0.227/0.272 | 0.218/0.284 |
| Number of non-hydrogen atoms | 19860 | 59580 | 5146 | 4980 |
| Protein residues | 2640 | 7920 | 660 | 660 |
| *R.m.s. deviations* |  |  |  |  |
| Bond lengths (Å) | 0.007 | 0.0142 | 0.01 | 0.004 |
| Bond angles (°) | 1.37 | 1.85 | 1.27 | 0.76 |
| *Ramachandran* |  |  |  |  |
| Favored (%) | 85.9 | 85.4 | 89.5 | 81.7 |
| Allowed (%) | 11.7 | 5.2 | 7.8 | 16.3 |
| Outliers (%) | 2.4 | 0.8 | 2.7 | 2.0 |
| Overall B | 101.5 | 106.0 | 60.3 | 120.7 |
| PDB ID | 7TJD | 7TJE | 7TJG | 7TJM1.85 |

## 3. Results

### 3.1. Qβ coat protein can assemble both T=1 and T=3 VLPs

Because the structure of a recombinantly expressed bacteriophage Qβ VLP has not been reported, we sought to determine the structure by X-ray crystallography. The recombinantly expressed and purified Qβ coat protein (amino acids 1-132) was crystallized, and the structure was determined. Interestingly, two forms of crystals are formed, with different morphologies. One form of crystal contains VLPs assembled in T=3 symmetry (Figure 1B), like previously observed (Golmohammadi et al., 1996) (Figure 1A). The most notable difference between this structure and previously reported structures is in particle size. The diameter of this particle is 166Å, much smaller than that of Qβ virus in the previously reported crystal (Golmohammadi et al., 1996) and cryo-EM (Cui et al., 2017; Gorzelnik et al., 2016) structures. The previously reported crystal structure (Golmohammadi et al., 1996)(PDB ID: 1QBE) has an asymmetric unit of 3 coat protein monomers that assemble a particle with 180 copies of the coat protein via crystallographic symmetry operations (Figure 1A). The first form of crystals reported here contain 60 copies of coat protein monomers in each asymmetric unit that assemble a particle with 180 copies of the coat protein via crystallographic symmetry operations (Figure 1B). However, the second form of crystals contain VLPs assembled in T=1 symmetry: each asymmetric unit contains 20 structurally independent chains of the coat protein, three asymmetric units assemble one VLP via a crystallographic 3-fold symmetry with 60 copies of the coat protein arranged in T=1 symmetry (Figure 1C).

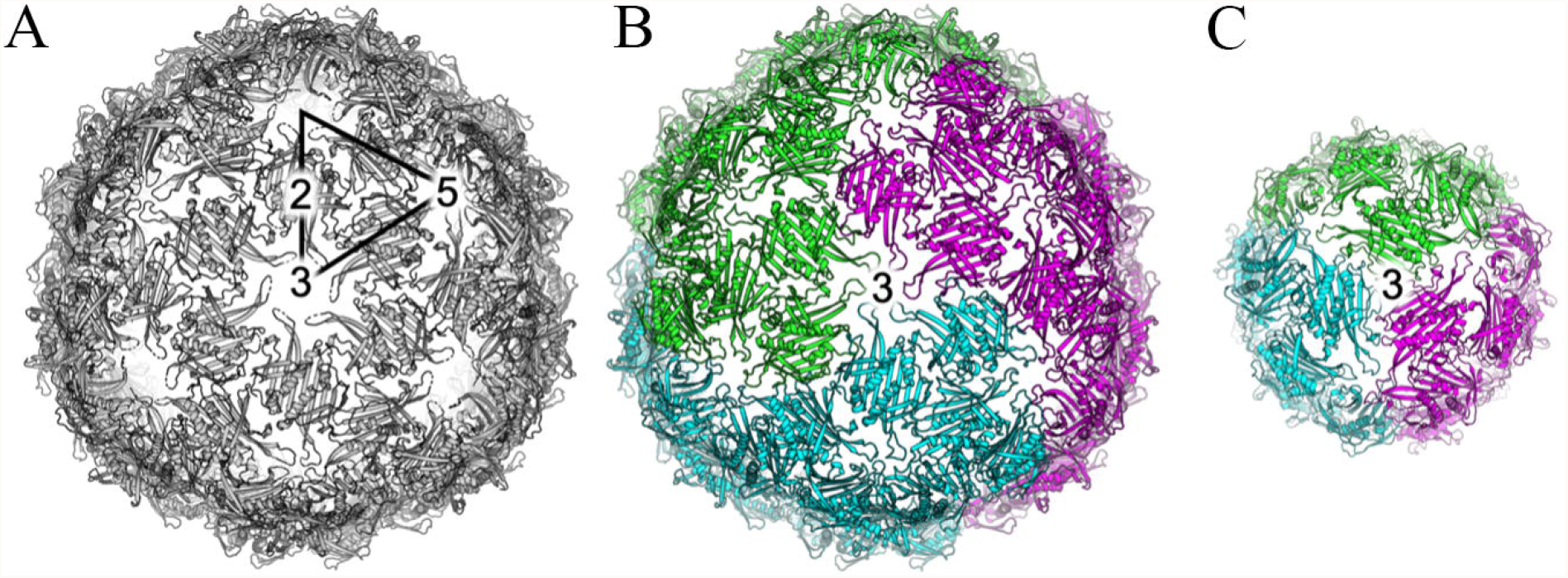

### 3.2. Comparison of Qβ coat protein monomers in various VLPs shows conformational flexibilities in inter-strand loops

As observed in the previous structure (PDB ID: 1QBE), each protomer is composed of seven beta strands followed by two alpha helices. The conformations of the 20 independent chains of our new T=1 structure and 60 independent chains of our new T=3 structure are similar to each other with the root mean square deviations (RMSDs) between chains range from 0.15 to 0.23 Å for the 132 Cα atoms. However, when compared to the structures of three independent chains of the previously reported crystal structure (PDB ID: 1QBE), apparent conformational differences in all of the inter-strand AB, BC, CD, DE, EF, and FG loops were observed (Figure 2). Of particular note, in the previous crystal structure (PDB ID: 1QBE), the residues 76-79 in the FG loop in chains A and C are at the five-fold vertices of the capsid particle, whereas the equivalent region in chain B at the six-fold vertices is missing. In our new structure, this segment, unambiguously resolved for all chains in the electron density map, participates only at the five-fold symmetry within a VLP. The discrepancy between the two VLP structures arises from the fact that in the T=3 structure, each monomer takes part in either a disulfide-linked pentamer or disulfide-linked hexamer (Figure 1B), whereas in our new T=1 structure, each monomer can only be part of a disulfide-linked pentamer (Figure 1C). The CD loop is observed at the three-fold vertices in both T=1 and T=3 particles, although their conformations are different (Figure 2).

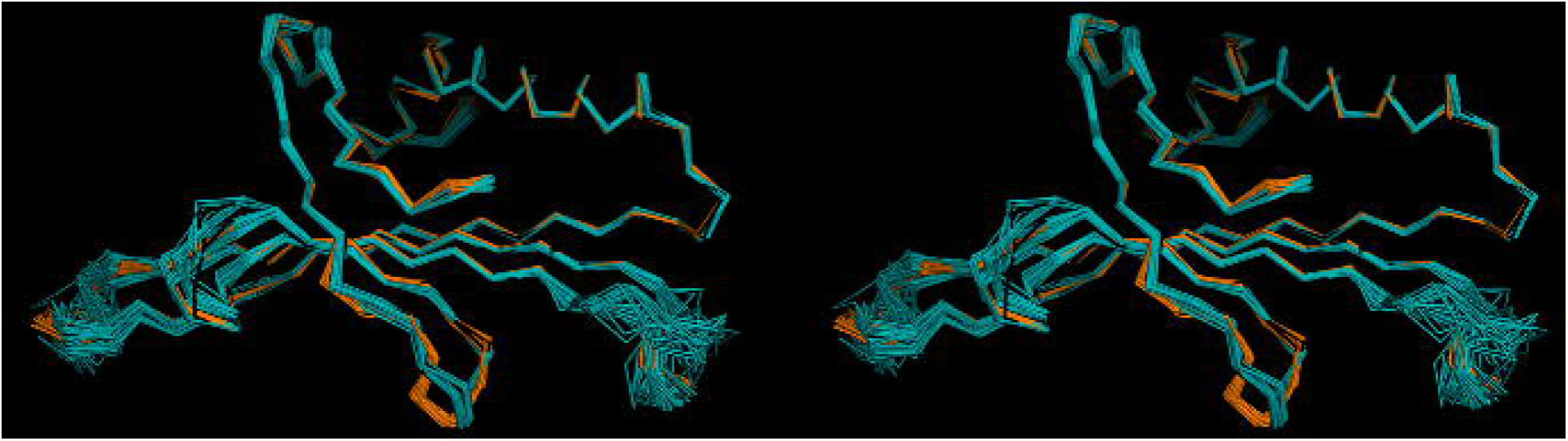

### 3.3. Qβ coat protein mutants also assemble VLPs in T=1 symmetry

To investigate the basis for the newly observed alternative assembly of Qβ VLPs, we determined the structures of two select mutants of Qβ coat proteins: A38K and A38K/A40C/D102C. The rationale for the mutant design has been reported previously (Sungsuwan, S. et al., 2021). Importantly, the crystals of both mutants are in space group I23 (Table 1), with five monomers in each asymmetric unit (Figure 3A). The five copies of coat protein monomers in each asymmetric unit assemble a particle containing 60 copies of the coat protein via crystallographic symmetry operations. This observation is consistent with the assembly mechanism of the new wild-type Qβ VLP in T=1 symmetry. This alternate symmetry does not appear to be due to the introduced mutations, as wild-type Qβ coat protein can also assemble T=1 VLPs. The comparison of three T=1 structures show no apparent conformational variations between wild-type and mutant VLPs except where mutations were introduced (Figure 3B).

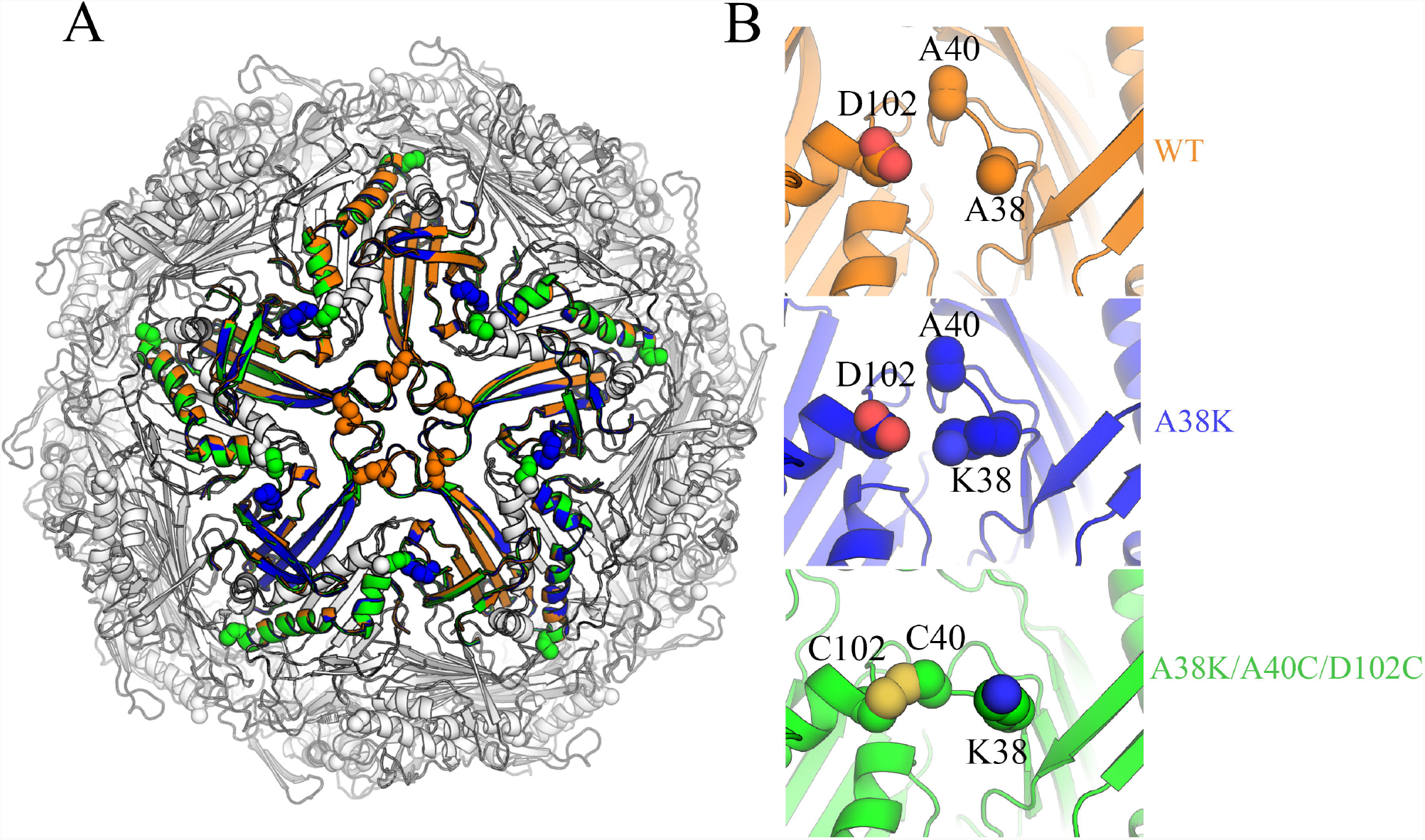

## 4. Discussion

The new crystal structures of Qβ VLPs we report here reveal that recombinantly expressed Qβ coat proteins can assemble VLPs in T=1 symmetry. The new structures of Qβ VLPs in T=1 symmetry are strikingly different from previously reported structures (Cui et al., 2017; Golmohammadi et al., 1996; Gorzelnik et al., 2016) of the Qβ virus in T=3 symmetry. This discrepancy is likely in part due to the fact the VLPs are formed by recombinant expression of the capsid protein in *E. coli* cells, while previous structural work used native virus samples. Our expression system lacks the genomic RNA of a native Qβ to template assembly. It was previously observed that the Qβ genomic RNA has many sites that can serve as “packaging signals” to interact with the coat protein to facilitate efficient and accurate virion assembly (Rolfsson et al., 2016). In the cryo-EM structure of native Qβ (Gorzelnik et al., 2016), numerous hairpins from the outmost layer of the genomic RNA make contacts with the coat protein, demonstrating the importance of genomic RNA in virion assembly. The lack of genomic RNA in our system is likely the driving force for arranging the coat protein in T=1 instead of T=3 symmetry in our new structure. In most other T=3 virus particles studied so far, chemically identical subunits have been observed to adapt to structurally different environments, causing problems for assembly. The assembly of most of these T=3 viruses is regulated by a folded “arm” present at some of the equivalent subunit interfaces, whereby this “arm” can behave like a switch to determine the type of contact and the local curvature of the particle shell. This “arm” appears to be essential for specification of the T=3 structure, because the coat protein subunits can form T=1 particles if this “arm” is removed (Erickson and Rossmann, 1982; Sorger et al., 1986). In Qβ VLPs, the dimer is relatively rigid and there are no equivalent “arms” to regulate the assembly. Instead, the FG loop can engage in both five-fold symmetry and six-fold symmetry in T=3 assembly but only in five-fold symmetry in a T=1 particle. In addition to the coat protein, the native virus capsid contains a readthrough version of the coat protein, called A1 (Takamatsu and Iso, 1982; Weber and Konigsberg, 1975), and a single monomer of the maturation protein, called A2 (Hohn and Hohn, 1970). Our expression system lacks the maturation protein (A2) found in native virus, which is necessary for infection (Hofstetter et al., 1974). However, in the cryo-EM structure (Gorzelnik et al., 2016), A2 appears to replace a coat protein dimer thereby breaking the coat symmetry, suggesting that lack of the A2 protein in our system does not underlie the T=1 assembly with the recombinantly produced coat protein.

Furthermore, our TEM data indicates that recombinantly expressed virus may be present in multiple sizes, which could include both T=1 and T=3 VLPs. We cannot rule out the possibility that VLPs are present in both assemblies in solution but only T=1 VLPs readily crystallize. Interestingly, a suspected T=1 Qβ VLP has been previously observed by TEM, among recombinantly expressed mutant (L35W) Qβ VLPs (Fiedler et al., 2012), however, this mutant also assembled in multiple intermediate sizes.

This intriguing observation sheds new light not only on the rational design of mutant Qβ VLPs but also on the physical principles that define the construction of viruses. Further research is needed to determine the effect of VLP size on trafficking to lymph nodes and B-cell engagement when used *in vivo* as a vaccine carrier.

## Acknowledgments

This research used resources of the Advanced Photon Source (APS), a U.S. Department of Energy (DOE) Office of Science User Facility operated for the DOE Office of Science by Argonne National Laboratory under Contract No. DE-AC02-06CH11357. Use of the LS-CAT Sector 21 was supported by the Michigan Economic Development Corporation and the Michigan Technology Tri-Corridor (Grant 085P1000817).

GM/CA at APS has been funded by the National Cancer Institute (ACB-12002) and the National Institute of General Medical Sciences (AGM-12006, P30GM138396). This research used resources of the Advanced Photon Source, a U.S. Department of Energy (DOE) Office of Science User Facility operated for the DOE Office of Science by Argonne National Laboratory under Contract No. DE-AC02-06CH11357. The Eiger 16M detector at GM/CA-XSD was funded by NIH grant S10 OD012289.

